# Fast-track adaptive laboratory evolution of *Cupriavidus necator* H16 with divalent metal cations

**DOI:** 10.1101/2023.10.24.563758

**Authors:** Sepwin Nosten Sitompul, Joseph Price, Kang Lan Tee, Tuck Seng Wong

## Abstract

Microbial strain improvement through adaptive laboratory evolution (ALE) has been a key strategy in biotechnology for enhancing desired phenotypic traits. Here, we present an accelerated ALE (aALE) workflow and its successful implementation in evolving *C. necator* H16 for enhanced tolerance towards elevated glycerol concentrations. The method involves the deliberate induction of genetic diversity through controlled exposure to divalent metal cations, enabling the rapid identification of improved variants. Through this approach, we observed the emergence of robust variants capable of growing in high glycerol concentration environments, demonstrating the efficacy of our aALE workflow. Our method offers several advantages over traditional ALE approaches, including its independence from genetically modified strains, specialized genetic tools, and potentially hazardous DNA-modifying agents. By utilizing divalent metal cations as mutagens, we offer a safer, more efficient, and cost-effective alternative for expansion of genetic diversity. With its ability to foster rapid microbial evolution, aALE serves as a valuable addition to the ALE toolbox, holding significant promise for the advancement of microbial strain engineering and bioprocess optimization.

## Introduction

Adaptive laboratory evolution (ALE) has evolved as an indispensable and robust technique, instrumental in enhancing a multitude of microbial properties vital for biomanufacturing [1-3]. These properties span from optimizing carbon utilization and improving microbial growth to enhancing tolerance against inhibitors and more. ALE stands out for its straightforward technical implementation and its applicability to diverse microbial strains. As a top-down methodology, it circumvents the need for intricate knowledge about the metabolic network of the microbe in question, as well as specialized equipment or a sophisticated genetic toolkit. This approach not only expedites the acquisition of valuable biological data but also facilitates reverse engineering, thereby unraveling crucial insights into the intricate dynamics of the phenotype-genotype interplay.

ALE has demonstrated its effectiveness in enhancing the properties of *Cupriavidus necator* H16, a betaproteobacterium renowned for its inherent ability to accumulate polyhydroxyalkanoate (PHA) up to 90% of its dry cell weight. PHA, a diverse range of thermoplastic polyesters, offers customizable material properties that can be fine-tuned through the manipulation of monomer composition and molecular weight. The applications of ALE on *C. necator* H16 are wide-ranging. Key examples include improving glycerol utilization [4], increasing halotolerance [5], improving growth on formate [6], and enhancing carbon monoxide tolerance [7].

Notwithstanding its simplicity, the ALE process can be time-consuming, involving serial or continuous cultivation under selective conditions. To expedite the evolutionary process, expanding genetic diversity becomes crucial, necessitating the creation of a larger and more complex pool of variants for subsequent selection. This is commonly achieved through the application of physical agents (*e*.*g*., UV radiation) and chemical agents (*e*.*g*., DNA alkylating agents like ethyl methanesulfonate). Although divalent metal ions, such as Mn^2+^, are extensively employed in random mutagenesis for protein evolution [8-9], their application in ALE remains unexplored.

This study seeks to introduce a novel, accelerated ALE workflow by incorporating divalent metal ions. Initially, the impact of three divalent metal ions (Co^2+^, Mn^2+^, and Zn^2+^) on the microbial growth of *C. necator* H16 was examined. Subsequently, these divalent metal ions were utilized to create genetic diversity and expedite the ALE process, facilitating the rapid identification of *C. necator* H16 variants capable of withstanding high concentrations of glycerol. This method marks the first application of divalent metal ions in ALE, presenting a significant advancement in the field.

## Material and Methods

### Bacterial strain and cultivation conditions

*C. necator* H16 (DSM 428) was cultivated in mineral salts medium (MSM; pH 7.0) with 1% (w/v) sodium gluconate [10] or nutrient broth (NB; 5 g/L peptone, 1 g/L beef extract, 2 g/L yeast extract, 5 g/L sodium chloride), supplemented with 10 μg/mL of gentamicin, at 30°C.

### The effect of divalent metal cations on the microbial growth of C. necator H16

To examine the impact of divalent metal cations on microbial growth, *C. necator* H16 was cultivated in Chi.Bio [11]. This process was carried out in a capped 30-mL clear borosilicate glass vial (Thermo Fisher Scientific, catalogue number 11593532), containing 15 mL of NB medium. The medium was supplemented with 10 μg/mL of gentamicin and varying concentrations of CoCl_2_ (0 – 20 μM), MnCl_2_ (0 – 25 mM), or ZnCl_2_ (0 – 5 mM). A fresh medium was inoculated with an overnight culture, targeting a starting OD_600_ of 0.2. The cultivation took place at a temperature of 30°C, with continuous stirring at a speed setting of 0.6 using a disc-shaped PTFE stir bar (Thermo Fisher Scientific, catalogue number 11878892). Optical density was measured at 650 nm at regular intervals during the experiment.

### Accelerated adaptive laboratory evolution (aALE) for high glycerol tolerance

The aALE process was conducted in accordance with the workflow presented in Figure 1. Each evolution cycle comprised two distinct stages: mutagenesis and selection. To initiate mutagenesis, *C. necator* H16 was cultured in a 50-mL Falcon tube, containing 5 mL of MSM with 1% (w/v) sodium gluconate. This medium was supplemented with 10 μg/mL of gentamicin and either 10 μM CoCl_2_, 15 mM MnCl_2_, or 0.7 mM ZnCl_2_, and allowed to incubate for 15 hours at 30°C. Subsequently, cells were collected through centrifugation at maximum speed for 2 minutes at room temperature. The cell pellet underwent two washes with 1 mL of MSM without any carbon source and was then appropriately diluted using the same medium. Moving to the selection stage, *C. necator* H16 was cultivated in Chi.Bio, utilizing a glass vial that contained 15 mL of MSM supplemented with 10% (v/v) glycerol, and 10 μg/mL of gentamicin. The medium was inoculated with the diluted cells obtained from the preceding mutagenesis step, targeting an initial OD_600_ of 0.2. Cultivation was sustained until an early stationary phase was achieved, thus signaling the commencement of a new evolution cycle.

**Figure 1:**
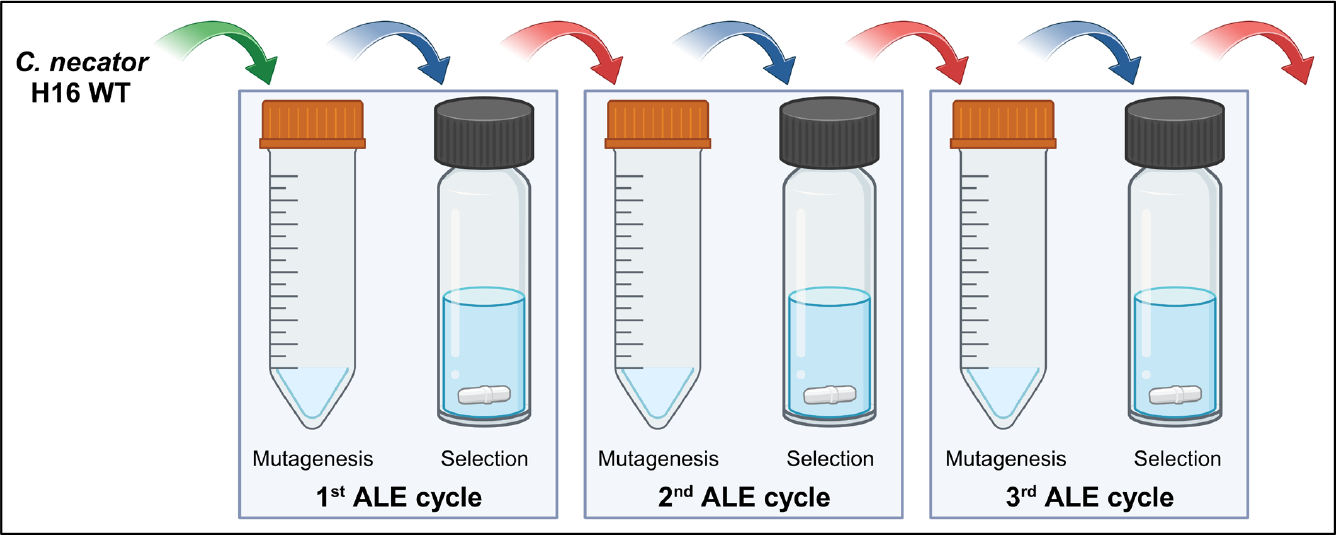
The workflow of accelerated adaptive laboratory evolution (aALE). Each aALE cycle involves the initial treatment of cells with 10 μM CoCl_2_, 15 mM MnCl_2_, or 0.7 mM ZnCl_2_ during the mutagenesis stage, followed by cultivation in selective medium during the selection stage.

### Adapted variants characterization in 96-well microplates and personal bioreactor

The adapted variants were subjected to characterization, with cultivation conducted either in a 96-well microplate or a personal bioreactor RST-1 (Biosan). For the 96-well microplate cultivation, each well was filled with 150 μL of MSM, supplemented with 10 μg/mL of gentamicin and 5% (v/v) glycerol. The microplate was placed on a Titramax 1000 shaker (Heidolph) operating at 30°C and 1050 rpm. Optical density was monitored at 595 nm using a Multiskan™ FC microplate photometer (Thermo Fisher Scientific). Alternatively, for cultivation in the personal bioreactor, a 50-mL TPP^®^ TubeSpin bioreactor tube with a membrane filter was employed. The tube was loaded with 20 mL of MSM supplemented with 10 μg/mL of gentamicin and 5 – 15% (v/v) glycerol. Cultivation took place at 30°C, with the tube rotated at a speed of 2000 rpm, incorporating a 1 s^-1^ reverse spin. Optical density was tracked at 850 nm during the cultivation process.

### Genomic DNA extraction

The extraction of genomic DNA was performed utilizing the GeneJET Genomic DNA Purification Kit (Thermo Fisher Scientific), following the manufacturer’s instructions. The eluted DNA underwent an additional purification step through ethanol precipitation. In a 1.5-mL microcentrifuge tube, 20 μL of 3 M sodium acetate (pH 5.2) was mixed with 200 μL of column-eluted DNA. Following this, 200 μL of ice-cold absolute ethanol was added and the resulting mixture was briefly vortexed. Subsequently, the tube was stored at -20°C overnight. After the overnight incubation, the tube was subjected to centrifugation at maximum speed for 30 minutes at 4°C. The DNA pellet was then washed twice with ice-cold 75% ethanol, followed by an incubation at 37°C for 15 – 20 minutes to remove any residual ethanol. The DNA pellet was finally reconstituted in an appropriate volume of Elution Buffer (10 mM Tris-Cl, pH 9.0, 0.1 mM EDTA) provided in the GeneJET kit. The concentration and purity of the DNA sample were determined using the EzDrop 1000 micro-volume spectrophotometer (Blue-Ray Biotech) and by electrophoresing 1 μg of DNA on a 1% (w/v) TAE gel.

### Genome sequencing

Microbial whole-genome sequencing was conducted by Novogene, employing the Illumina PE150 technology. Subsequently, the sequencing data was aligned with the reference sequences (GenBank CP039287.1, CP039288.1, and CP039289.1) through the use of BWA [12]. Detection of single nucleotide variations (SNVs) and insertions/deletions (InDels) was executed using the GATK software [13]. Variation annotation was performed utilizing the capabilities of ANNOVAR [14].

## Results and Discussion

### Divalent metal cations affect the microbial growth of C. necator H16

Metal ions play a vital role in sustaining life, being essential for bacterial growth, with Zn^2+^, Mn^2+^, and Fe^2+^ cellular concentrations falling within the range of 0.4 – 1 mM under favorable conditions [15]. Metalloproteins depend on the presence of these ions to execute crucial functions, contributing to both the structure of proteins and the facilitation of catalytic activities [16]. Instances of bacterial growth inhibition due to metal intoxication can result from the generation of harmful reactive oxygen species or the erroneous metallation of enzymes pivotal in critical metabolic pathways. Additionally, surpassing the threshold concentrations of these metal ions can disrupt the accuracy of DNA synthesis [17-18].

In this study, CoCl_2_, MnCl_2_, and ZnCl_2_ were specifically chosen to evaluate their effects on microbial growth and their potential as chemical mutagens, intended to enhance the genetic diversity for ALE. CoCl_2_ and MnCl_2_ were categorized as class 1 compounds by Loeb and colleagues, leading to an elevation in error frequency and a decline in DNA synthesis *in vitro* [17]. Conversely, ZnCl_2_ was classified as a class 2 compound, which did not compromise fidelity but did result in reduced DNA synthesis. Notably, the mutagenic impact of 4 mM CoCl_2_ and 10 mM MnCl_2_ was observed to reduce fidelity by a minimum of 30%, while 0.4 mM ZnCl_2_ had no significant effect on fidelity [17]. Furthermore, MnCl_2_ is frequently utilized in error-prone polymerase chain reaction (epPCR) for protein evolution, typically within the concentration range of 0.01 – 0.05 mM [19-20].

Among the three metal salts studied, *C. necator* H16 exhibited the highest susceptibility to CoCl_2_ (Figure 2). At concentrations of 8 μM and 10 μM of CoCl_2_, a significant reduction in growth was observed, with concentrations exceeding 10 μM proving to be severely detrimental to growth. In contrast, *C. necator* H16 demonstrated a higher tolerance for MnCl_2_, as concentrations up to 10 mM had no significant impact on microbial growth. Even at concentrations as high as 15 mM and 25 mM, growth was observed, albeit at a slower rate. Addition of 0.6 – 0.8 mM ZnCl_2_ led to a reduced growth, with minimal growth observed beyond this concentration range. Based on the growth data, the concentrations of 10 μM CoCl_2_, 15 mM MnCl_2_, and 0.7 mM ZnCl_2_ were selected to establish an aALE workflow. At these specific concentrations, microbial growth was feasible, although at a reduced rate, potentially indicating a decrease in DNA synthesis and/or the introduction of errors during DNA replication. The latter effect was sought to increase genetic diversity.

**Figure 2:**
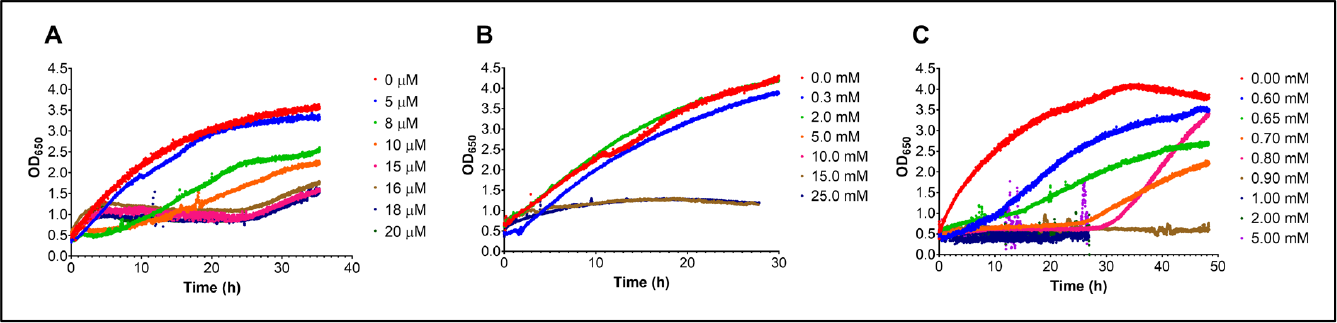
The influence of divalent metal cations at different concentrations, 0 – 20 μM CoCl_2_ (**A**), 0 – 25 mM MnCl_2_ (**B**), and 0 – 5 mM ZnCl_2_ (**C**), on the microbial growth of *C. necator* H16.

### Divalent metal ions altered the growth performance of microbial populations in 10% (v/v) glycerol

To expedite the process of ALE, we adopted a scheme commonly employed for protein evolution, which involves creating genetic diversity followed by selecting improved variants. In our aALE workflow (Figure 1), cells were initially subjected to random mutagenesis through cultivation in the presence of 10 μM CoCl_2_, 15 mM MnCl_2_, or 0.7 mM ZnCl_2_. Subsequently, the cells were cultivated under selective conditions to enrich the population displaying the desired phenotype.

In order to showcase the practical application of the aALE method, we opted to adapt *C. necator* H16 to utilize 10% (v/v) glycerol. This particular adaptation was guided by two key considerations. Firstly, glycerol stands as an appealing feedstock for biomanufacturing purposes [21-22]. Secondly, while ALE has been applied to enhance glycerol utilization in various microbes, it has typically been carried out at lower glycerol concentrations. For example, adaptation of *C. necator* H16 was conducted at 0.5% (v/v) glycerol [4], *E. coli* at 0.2% (v/v) [23], and *S. cerevisiae* at 1% (v/v) [24]. The adaptation to 10% (v/v) glycerol is challenging, likely due to the cellular dehydration experienced, which adversely affects cell viability [25].

Through the application of aALE, we successfully generated *C. necator* H16 microbial populations that exhibited a higher growth rate and reached higher final OD_650_ values in the presence of 10% (v/v) glycerol (as indicated by the arrows in Figure 3). This effect was noticeable after 3 cycles of evolution when cells were exposed to 10 μM CoCl_2_, with each cycle defined as a sequence of mutagenesis followed by selection. When cells were treated with 15 mM MnCl_2_ and 0.7 mM ZnCl_2_, this enhanced growth was observed after only 2 cycles of evolution. Continued iterations revealed an opposing trend, characterized by slower growth and lower final OD_650_ values, which suggested the over-accumulation of detrimental mutations.

**Figure 3:**
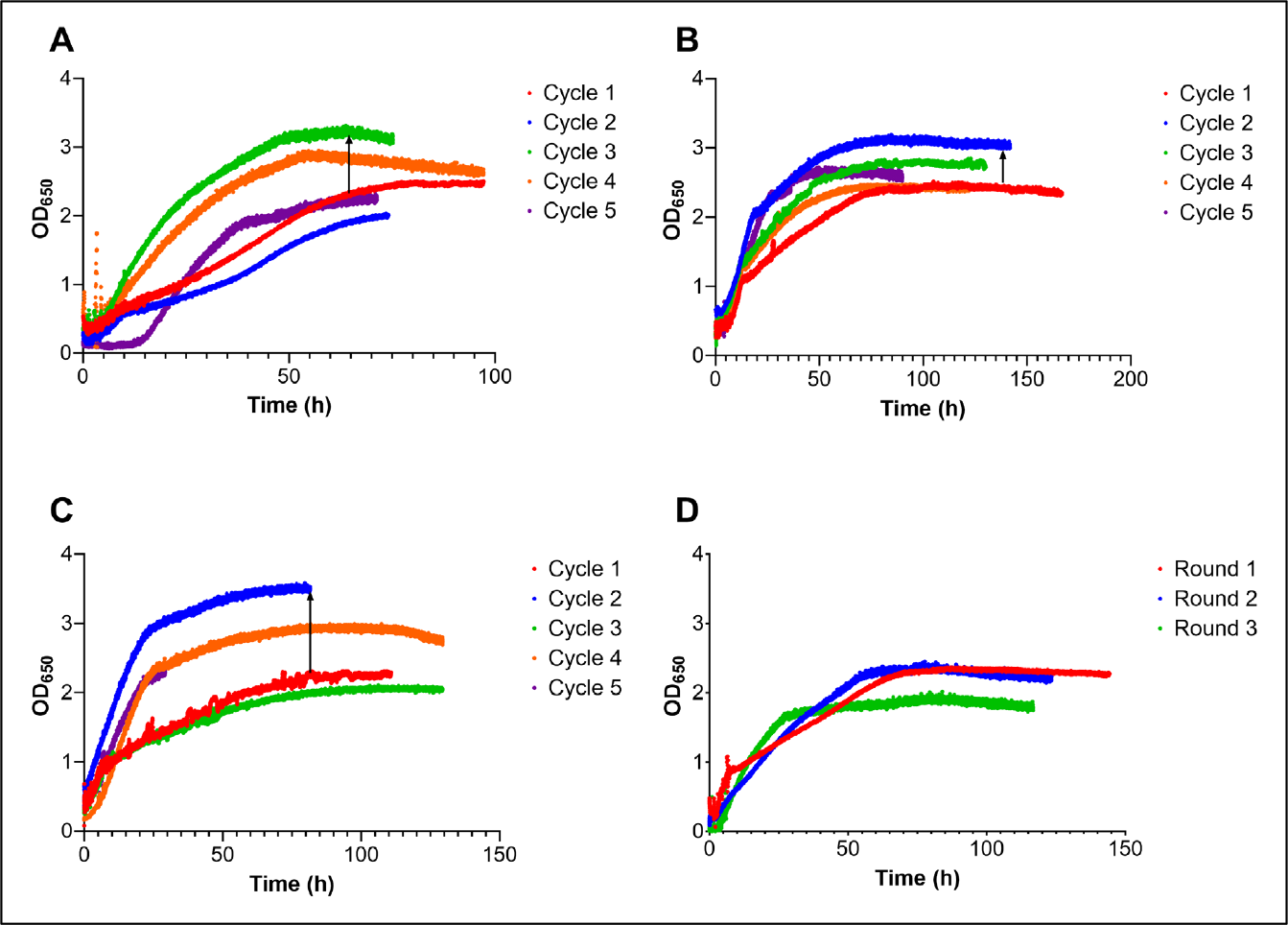
The growth of *C. necator* H16 in 10% (v/v) glycerol in Chi.Bio, following treatment with 10 μM CoCl_2_ (**A**), 15 mM MnCl_2_ (**B**), 0.7 mM ZnCl_2_ (**C**), and in the absence of treatment (**D**). The black arrows signify an increase in the final OD_650_ values.

Furthermore, we conducted the cultivation of *C. necator* H16 in 10% (v/v) glycerol without any prior treatment involving divalent metal cations (Figure 3D). It is interesting to note that *C. necator* H16 demonstrated the capacity to grow at this concentration. However, we observed a gradual reduction in growth during subsequent cultivation rounds, notably reflected in a decrease in the final OD_650_ value. This implies that achieving an improved performing population would require an extended duration if we were to continue with successive cultivation in 10% (v/v) glycerol. Leveraging the amplification of genetic diversity facilitated by the use of divalent metal cations, we captured rare beneficial mutations, effectively shortening the time required for enhancement.

### Adapted variants grew better at 10% (v/v) and 15% (v/v) glycerol

Populations from the second cycle (MnCl_2_- or ZnCl_2_-treated cells) and the third cycle (CoCl_2_-treated cells) were spread on an agar plate, with subsequent individual positive clones subjected to further confirmation and detailed characterization.

In Figure 4, we compared the growth performance of *C. necator* H16 variants (Mn-C2-B11 and Mn-C2-F11), isolated from the population that underwent 2 cycles of 15 mM MnCl_2_ treatment, against the wild type (WT) and v6C6 variant obtained from a prior ALE study [4]. The v6C6 variant was isolated after 6 rounds of sequential cultivation at 0.5% (v/v) glycerol. As part of this comparison, we included a variant (Mn-C1-D3) isolated from the population that underwent only 1 cycle of 15 mM MnCl_2_ treatment, aiming to assess whether a single aALE cycle sufficed to isolate improved variants. Growth comparisons were conducted in 96-well microplate using two carbon sources: 1% (w/v) sodium gluconate, a preferred carbon source of *C. necator* H16 [26], and 5% (v/v) glycerol. All variants exhibited similar growth in gluconate.

**Figure 4:**
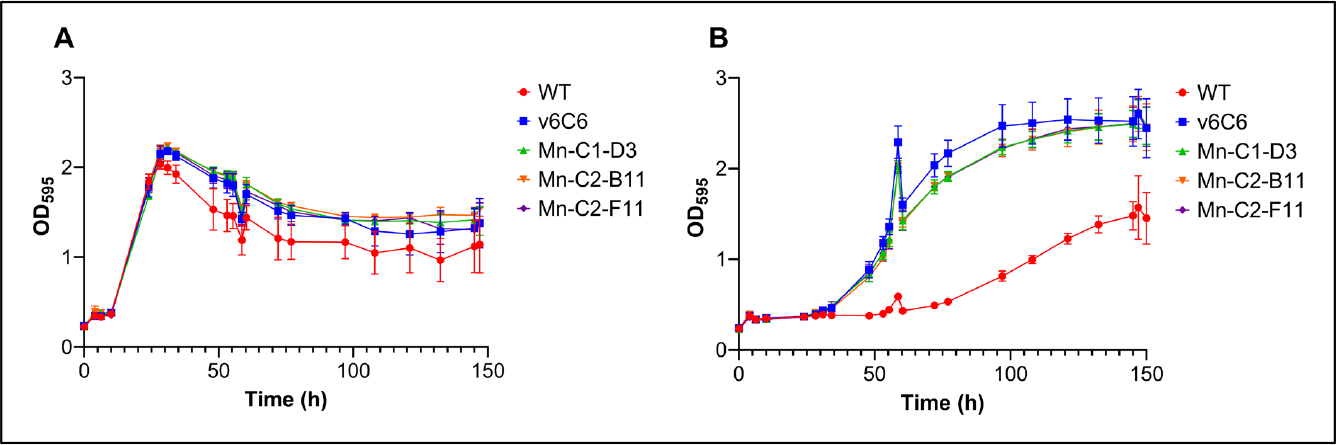
The growth of variants Mn-C1-D3 (green), Mn-C2-B11 (orange), and Mn-C2-F11 (purple) in comparison to the WT (red) and v6C6 (blue), in 1% (w/v) sodium gluconate (**A**) and 5% (v/v) glycerol (**B**). The cultivation was conducted in a 96-well microplate. Variant Mn-C1-D3 was obtained from the population after 1 cycle of aALE, while variants Mn-C2-B11 and Mn-C2-F11 were isolated after 2 cycles of aALE. Cultivations were performed in triplicate, and the error bars indicate standard deviations.

However, at 5% (v/v) glycerol, all ALE-derived variants demonstrated superior growth compared to the wild type. The v6C6 variant outperformed all other variants at this particular glycerol concentration. Remarkably, the performance of the Mn-C1-D3 variant was found to be on par with that of v6C6, underscoring the advantage of aALE in expediting the process of identifying improved variants. Furthermore, we characterized the variants derived from populations treated with CoCl_2_ and ZnCl_2_ and noted similar trends (Figure S1).

Subsequently, we conducted a comprehensive growth comparison involving variant Mn-C2-B11 alongside the WT, v6C6, Co-C3-F3, and Zn-C2-G5, at a larger cultivation scale of 20 mL in three distinct glycerol concentrations [5% (v/v), 10% (v/v), and 15% (v/v)], employing a personal bioreactor (Figure 5). Variants Co-C3-F3 and Zn-C2-G5 were isolated from populations that underwent 3 cycles of 10 μM CoCl_2_ treatment and 2 cycles of 0.7 mM ZnCl_2_ treatment, respectively. At 5% (v/v) glycerol, the results were consistent with the findings obtained earlier using the 96-well microplate. v6C6 remained the most proficient variant at this concentration, closely trailed by Mn-C2-B11. However, at 10% (v/v) glycerol, the concentration at which aALE was conducted, all aALE variants exhibited improved growth compared to v6C6. Surprisingly, v6C6 exhibited slower growth compared to the WT at this concentration. Additionally, we conducted a comparative growth at 15% (v/v), a significantly higher concentration than the aALE condition of 10% (v/v). The results demonstrated a consistent trend, with Mn-C2-B11 emerging as the superior performer at this concentration. The disparities observed between the 5% (v/v) and higher concentrations [10% (v/v) or 15% (v/v)] potentially underscored the superior ability of variants obtained at 10% (v/v) to withstand high osmotic stress, in addition to displaying improved glycerol utilization.

**Figure 5:**
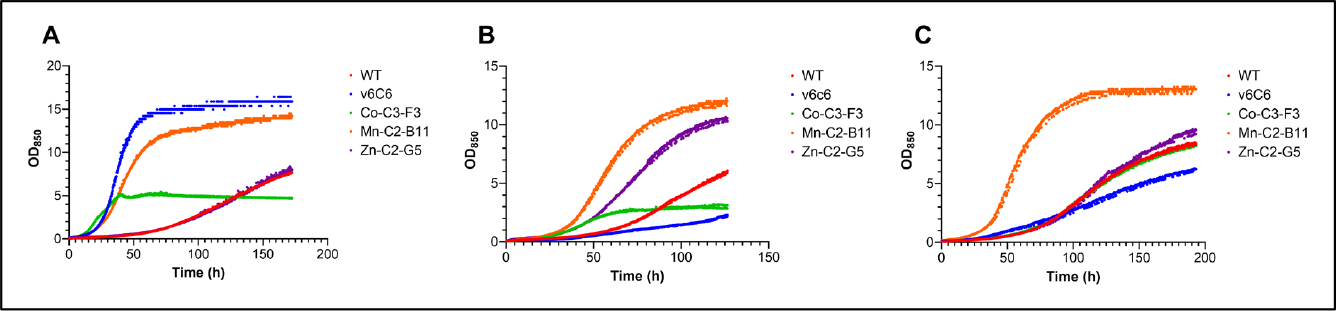
The growth of variants Co-C3-F3 (green), Mn-C2-B11 (orange), and Zn-C2-G5 (purple), in comparison to the WT (red) and v6C6 (blue), in 5% (v/v) glycerol (**A**), 10% (v/v) glycerol (**B**), and 15% (v/v) glycerol (**C**). The cultivations were carried out in a personal bioreactor.

### SNVs and InDels were detected in all aALE variants

The genomes of the WT, Mn-C2-B11, Co-C3-F3, and Zn-C2-G5 were sequenced and subsequently compared against the reference genomes (GenBank accession numbers CP039287, CP039288, and CP039289) [27]. Single nucleotide variations (SNVs) (supplementary Table S1) and insertions/deletions (InDels) (supplementary Table S2) were detected in all variants originating from aALE. These mutations are primarily located in chromosome 1. MnCl_2_ and CoCl_2_ exhibited significantly higher mutagenic effects compared to ZnCl_2_ (supplementary Table S3), in line with the findings previously documented by Loeb and colleagues [17-18].

Typically transition-heavy mutations are generated in epPCR [8-9]. In our aALE-derived variants, we identified both transition and transversion mutations. Moreover, the observed mutation count was higher in comparison to the variants obtained through serial cultivation, such as v6C6 [4].

The dataset from the small number of genome sequences was limited and insufficient to draw a broad conclusion. Nevertheless, our sequencing results confirmed two significant aspects: the induction of mutations by divalent metal cations and their effective application in expanding genetic diversity.

### Variant Mn-C2-B11 carries two nonsynonymous mutations in the gene encoding glycerol kinase

Variant Mn-C2-B11, displaying the most robust growth at 10% (v/v) and 15% (v/v) glycerol (Figure 5), carries two nonsynonymous mutations in the *glpK* gene, resulting in A264V and W480S substitutions within the glycerol kinase (GlpK) sequence. The occurrence of two mutations within a single protein-coding gene is statistically uncommon and underscores the impact of treating cells with 15 mM MnCl_2_, effectively inducing a high mutation rate that aligns precisely with our intended outcomes within the aALE workflow. GlpK plays a crucial role in the initial step of glycerol metabolism, facilitating the conversion of triol into glycerol-3-phosphate. Moreover, the W480S substitution in GlpK was also observed in the Co-C3-F3 variant within this study and the v6C6 variant from a previous ALE study [4]. Additionally, mutations in the *glpK* gene were also highly recurrent in a glycerol-adapted *E. coli* [23, 28-31] (supplementary Table S4), emphasizing the significance of GlpK in glycerol assimilation.

While the functional consequence of the W480S substitution remains unclear, it is worth noting that residue A264 is positioned within the active site of GlpK, adjacent to D261 and Q262, which are highly conserved among glycerol kinase sequences and play a role in interacting with glycerol [32] (Figure S2). Based on this context, we firmly believe that the A264V substitution modifies substrate interactions. A comprehensive biochemical characterization of GlpK variants would be essential to gain a thorough understanding of the functional implications of these mutations. However, this analysis falls outside the scope of this article, which centers on the aALE workflow.

### Potential augmented stress tolerance in aALE variants

Variants Mn-C2-B11 and Co-C3-F3 also carry a G253W substitution in the YgcG family protein, a protein that is predicted to have a transmembrane domain and is considered a putative phosphatase. This substitution was absent in variant Zn-C2-G5 and the WT, which served as the starting point for our aALE (supplementary Table S1). Examination using STRING [33] indicated that YgcG is functionally associated with the membrane stress resistance protein YqcG in *E. coli* (Figure S3). Intriguingly, the same protein was mutated at the identical position in a recent ALE study, where *C. necator* H16 underwent evolution for enhanced halotolerance [5]. Commencing from a W253 genotype, the identified halotolerant variant carried a W253G mutation, thereby reversing W253 back to G253.

## Conclusions

In conclusion, we have successfully devised an accelerated and cost-effective ALE workflow, demonstrating its effectiveness in evolving *C. necator* H16 for enhanced tolerance towards high glycerol concentrations. This method offers several advantages. Firstly, by amplifying genetic diversity, we significantly reduce the time required to attain a desired phenotype. In this study, we were able to identify an improved variant after just 1 cycle of aALE, underscoring the efficiency of this approach. By way of comparison, a previous study on ALE of *E. coli* MG1655 in M9 minimal medium supplemented with 0.2% glycerol showed that growth rates could be increased by more than 2-fold through log-phase serial transfers spanning over 800 generations [31]. In another related work, *E. coli* W strain underwent adaptive evolution over 1300 generations [23]. Secondly, our aALE workflow does not necessitate a genetically modified strain (*e*.*g*., the use of a *mutS* knock-out derivative [31]) to initiate an ALE, nor does it rely on specialized genetic tools (*e*.*g*., Tn5 mutagenesis [34]). Thirdly, our method does not depend on DNA-modifying agents, which are frequently categorized as carcinogens. Lastly, this methodology can be readily adapted to other microbial strains, enhancing its versatility and applicability.

## Acknowledgement

We express our sincere gratitude to the following organizations for their financial support: the UKRI Global Challenges Research Fellowship (to KLT), the RAEng | Leverhulme Trust Senior Research Fellowship (LTSRF1819\15\21, to TSW), the National Research Council of Thailand (P2250317/3, to TSW), and the Directorate General of Higher Education, Research, and Technology, Ministry of Education, Culture, Research, and Technology of the Republic of Indonesia (PhD scholarship to SNS).

