## Supplementary Information for "Fast-track adaptive laboratory evolution of *Cupriavidus necator* H16 with divalent metal cations"

[1]Department of Chemical & Biological Engineering, University of Sheffield, Sir Robert Hadfield Building, Mappin Street, Sheffield S1 3JD, United Kingdom; [2]Evolutor Ltd, The Innovation Centre, 217 Portobello, Sheffield S1 4DP, United Kingdom; [3]National Center for Genetic Engineering and Biotechnology (BIOTEC), National Science & Technology Development Agency (NSTDA), 113 Thailand Science Park, Phahonyothin Road, Khlong Nueng, Khlong Luang, Pathum Thani 12120, Thailand; [4]School of Pharmacy, Bandung Institute of Technology, Bandung, West Java, Indonesia.

***Corresponding authors:**

**Dr. Kang Lan Tee**


**Prof. Tuck Seng Wong**


**Table S1:** SNVs detected in Co-C3-F3, Mn-C2-B11 and Zn-C2-G5 are highlighted in yellow (**US**: upstream, **DS**: downstream)

| Chromosome | Location | Position | Locus tag | Gene | WT | | Co-C3-F3 | | Mn-C2-B11 | | Zn-C2-G5 | |
|---|---|---|---|---|---|---|---|---|---|---|---|---|
| | | | | | REF | ALT | REF | ALT | REF | ALT | REF | ALT |
| NZ_CP039287.1 | US/DS | 192516 | gene-E6A55_RS00915;gene-E6A55_RS00905,gene-E6A55_RS00910 | | G | T | G | T | | | G | T |
| NZ_CP039287.1 | US/DS | 333421 | gene-E6A55_RS01595;gene-E6A55_RS01590 | | G | T | | | G | T | G | T |
| NZ_CP039287.1 | US/DS | 333427 | gene-E6A55_RS01595;gene-E6A55_RS01590 | | G | T | | | G | T | G | T |
| NZ_CP039287.1 | Exonic | 500511 | gene-E6A55_RS02390 | amino acid ABC transporter permease | G | A | | | | | G | A |
| NZ_CP039287.1 | Exonic | 572008 | gene-E6A55_RS02750 | recombinase RecA | | | C | T | C | T | | |
| NZ_CP039287.1 | Exonic | 578522 | gene-E6A55_RS02785 | ligase | | | A | G | A | G | | |
| NZ_CP039287.1 | Exonic | 738904 | gene-E6A55_RS03485 | ornithine cyclodeaminase | | | T | A | T | A | | |
| NZ_CP039287.1 | Exonic | 791503 | gene-E6A55_RS03700 | YgcG family protein | | | C | A | C | A | | |
| NZ_CP039287.1 | US/DS | 1166481 | gene-E6A55_RS05420;gene-E6A55_RS05410,gene-E6A55_RS05415 | | T | C | | | | | T | C |
| NZ_CP039287.1 | Exonic | 1485292 | gene-E6A55_RS06940 | PAS domain-containing sensor histidine kinase | | | G | C | G | C | | |
| NZ_CP039287.1 | ncRNA-exonic | 1807929 | rna-E6A55_RS08390 | | | | A | T | | | | |
| NZ_CP039287.1 | ncRNA-exonic | 1807930 | rna-E6A55_RS08390 | | | | T | A | | | | |
| NZ_CP039287.1 | Exonic | 2101221 | gene-E6A55_RS09900 | D-galactonate dehydratase family protein | | | A | C | | | | |
| NZ_CP039287.1 | Exonic | 2597061 | gene-E6A55_RS12270 | adenosylcobalamin-dependent ribonucleoside-diphosphate reductase | C | A | C | A | C | A | | |
| NZ_CP039287.1 | US/DS | 2612983 | rna-E6A55_RS12325,rna-E6A55_RS12330;gene-E6A55_RS12320,rna-E6A55_RS12335,rna-E6A55_RS12340,rna-E6A55_RS12345,rna-E6A55_RS12350 | | | | | | T | C | | |
| NZ_CP039287.1 | US/DS | 2612986 | rna-E6A55_RS12325,rna-E6A55_RS12330;gene-E6A55_RS12320,gene-E6A55_RS12355,rna-E6A55_RS12335,rna-E6A55_RS12340,rna-E6A55_RS12345,rna-E6A55_RS12350 | | | | | | A | T | | |
| NZ_CP039287.1 | US/DS | 2612991 | rna-E6A55_RS12325,rna-E6A55_RS12330;gene-E6A55_RS12320,gene-E6A55_RS12355,rna-E6A55_RS12335,rna-E6A55_RS12340,rna-E6A55_RS12345,rna-E6A55_RS12350 | | | | | | G | A | | |
| NZ_CP039287.1 | US/DS | 2626787 | gene-E6A55_RS12395,gene-E6A55_RS34550,gene-E6A55_RS35235;gene-E6A55_RS12400,gene-E6A55_RS12405,gene-E6A55_RS12410 | | | | | | C | G | | |
| NZ_CP039287.1 | US/DS | 2626790 | gene-E6A55_RS12395,gene-E6A55_RS34550,gene-E6A55_RS35235;gene-E6A55_RS12400,gene-E6A55_RS12405,gene-E6A55_RS12410 | | | | | | T | G | | |
| NZ_CP039287.1 | US/DS | 2626815 | gene-E6A55_RS12395,gene-E6A55_RS34550,gene-E6A55_RS35235;gene-E6A55_RS12400,gene-E6A55_RS12405,gene-E6A55_RS12410 | | | | | | A | C | | |
| NZ_CP039287.1 | Exonic | 2721992 | gene-E6A55_RS12840 | glycerol kinase | | | | | C | T | | |
| NZ_CP039287.1 | Exonic | 2722640 | gene-E6A55_RS12840 | glycerol kinase | | | G | C | G | C | | |
| NZ_CP039287.1 | Exonic | 2768732 | gene-E6A55_RS13105 | RseB | | | | | | | G | C |
| NZ_CP039287.1 | Exonic | 3718338 | gene-E6A55_RS17570 | penicillin-binding protein 1A | G | T | | | | | G | T |
| NZ_CP039287.1 | Exonic | 3981763 | gene-E6A55_RS18850 | response regulator | C | T | | | | | C | T |
| NZ_CP039288.1 | Exonic | 56691 | gene-E6A55_RS19345,gene-E6A55_RS19350 | MFS transporter | | | A | C | | | | |
| NZ_CP039288.1 | Exonic | 56695 | gene-E6A55_RS19345,gene-E6A55_RS19350 | MFS transporter | | | | | | | A | C |
| NZ_CP039288.1 | Exonic | 164765 | gene-E6A55_RS19830 | sigma-54-dependent Fis family transcriptional regulator | C | T | | | | | C | T |
| NZ_CP039288.1 | ncRNA-exonic | 177502 | rna-E6A55_RS19890 | | | | T | G | | | | |
| NZ_CP039288.1 | Exonic | 649721 | gene-E6A55_RS22090 | hypothetical protein | | | G | C | G | C | | |
| NZ_CP039288.1 | Exonic | 1205170 | gene-E6A55_RS24600 | MurR/RpiR family transcriptional regulator | | | C | T | C | T | | |
| NZ_CP039288.1 | Exonic | 1628671 | gene-E6A55_RS26535 | formate dehydrogenase-N subunit alpha | T | C | | | | | T | C |
| NZ_CP039288.1 | Exonic | 1820152 | gene-E6A55_RS27345 | VWA domain-containing protein | | | G | C | G | C | | |
| NZ_CP039288.1 | Exonic | 2496322 | gene-E6A55_RS30325 | tetratricopeptide repeat protein | | | | | | | G | T |
| NZ_CP039289.1 | Exonic | 682 | gene-E6A55_RS32220 | twin-arginine translocation signal domain-containing protein | C | A | | | | | C | A |
| NZ_CP039289.1 | Exonic | 202180 | gene-E6A55_RS33140 | nucleotidyl transferase AbiEii/AbiGii toxin family protein | | | T | C | T | C | | |

**Table S2:** InDels detected in Co-C3-F3, Mn-C2-B11 and Zn-C2-G5 are highlighted in yellow (**US**: upstream, **DS**: downstream)

| Chromosome | Location | Position 1 | Position 2 | Locus tag | Function | WT | | Co-C3-F3 | | Mn-C2-B11 | | Zn-C2-G5 | |
|---|---|---|---|---|---|---|---|---|---|---|---|---|---|
| | | | | | | REF | ALT | REF | ALT | REF | ALT | REF | ALT |
| NZ_CP039287.1 | Exonic | 1309779 | 1309779 | gene-E6A55_RS34760 | Hypothetical protein | - | CAGCAG CCGGAG CAG | | | | | | |
| NZ_CP039287.1 | Exonic | 1712937 | 1712984 | gene-E6A55_RS07985 | DNA segregation ATPase FtsK/SpoIIIE or related protein | AGCCGA AGCCGA AGCCGA AGCCGA AGCCGA AGCCGA AGCCGA AGCCGA | - | | | | | AGCCGA AGCCGA AGCCGA AGCCGA AGCCGA AGCCGA AGCCGA AGCCGA | - |
| NZ_CP039287.1 | ncRNA_exonic | 1807932 | 1807932 | rna-E6A55_RS08390 | 23S rRNA | | | - | GCAC | | | | |
| NZ_CP039287.1 | US/DS | 2626802 | 2626805 | gene-E6A55_RS12395,gene-E6A55_RS34550,gene-E6A55_RS35235;gene-E6A55_RS12400,gene-E6A55_RS12405,gene-E6A55_RS12410 | Predicted arabinose efflux permease AraJ, MFS family | | | | | TGTG | - | | |
| NZ_CP039287.1 | US/DS | 2626808 | 2626810 | gene-E6A55_RS12395,gene-E6A55_RS34550,gene-E6A55_RS35235;gene-E6A55_RS12400,gene-E6A55_RS12405,gene-E6A55_RS12410 | Predicted arabinose efflux permease AraJ, MFS family | | | | | TGG | - | | |
| NZ_CP039287.1 | Exonic | 3720874 | 3720877 | gene-E6A55_RS34910 | Hypothetical protein | | | GCCA | - | | | | |
| NZ_CP039287.1 | Exonic | 3720882 | 3720883 | gene-E6A55_RS34910 | Hypothetical protein | | | TG | - | | | | |
| NZ_CP039287.1 | Exonic | 3800472 | 3800472 | gene-E6A55_RS18020 | DNA-binding transcriptional regulator, Lrp family | | | G | - | G | - | | |
| NZ_CP039287.1 | Exonic | 3943689 | 3943700 | gene-E6A55_RS18695 | Choline dehydrogenase or related flavoprotein | | | CAGGCG GTTGGC | - | CAGGCG GTTGGC | - | | |
| NZ_CP039288.1 | ncRNA_exonic | 177504 | 177504 | rna-E6A55_RS19890 | 23S rRNA | | | - | G | | | | |
| NZ_CP039288.1 | Exonic | 265365 | 265365 | gene-E6A55_RS20305 | Flagellar motor component MotA | - | GGCGCC GCCTTC | | | | | - | GGCGCC GCCTTC |
| NZ_CP039288.1 | Exonic | 649714 | 649714 | gene-E6A55_RS22090 | Hypothetical protein | | | | | - | CCCCCC | | |
| NZ_CP039288.1 | Exonic | 2219351 | 2219372 | gene-E6A55_RS29140 | Membrane carboxypeptidase/penicillin-binding protein | | | GCGCCG GCGTGC GCCGCT ACCT | - | GCGCCG GCGTGC GCCGCT ACCT | - | | |
| NZ_CP039288.1 | Exonic | 2459882 | 2459882 | gene-E6A55_RS30165 | Glycosyltransferase, catalytic subunit of cellulose synthase and poly-beta-1,6-N-acetylglucosamine synthase | | | | | - | CGCCCC C | | |
| NZ_CP039289.1 | Exonic | 204769 | 204769 | gene-E6A55_RS33155 | HAMP domain-containing histidine kinase | | | - | CGGGCT | | | | |

**Table S3:** Analysis of mutational spectra

| Nucleotide substitution | Co-C3-F3 | Mn-C2-B11 | Zn-C2-G5 |
|---|---|---|---|
| $T_s$, A→G, T→C | 5 | 6 | 0 |
| $T_s$, G→A, C→T | 4 | 6 | 0 |
| $T_v$, A→T, T→A | 3 | 2 | 0 |
| $T_v$. A→C, T→G | 7 | 5 | 2 |
| $T_v$, G→C, C→G | 4 | 5 | 1 |
| $T_v$, G→T, C→A | 1 | 1 | 1 |
| $T_s/T_v$ | 0.6 | 0.9 | 0 |
| InDel | 11 | 10 | 1 |

**Table S4:** Mutations found in the glycerol kinase sequences of glycerol-adapted microbes through ALE

| Organism | Amino acid substitution* | Possible functional consequence | Reference |
|---|---|---|---|
| *C. necator* H16 | A264V | Altered interaction with glycerol | This study |
| *C. necator* H16 | W480S | Unknown | This study and [1] |
| *Escherichia coli* W | T236M | Reduced or diminished sensitivity towards inhibitory effects of fructose 1,6-bisphosphate (FBP) | [2] |
| *Escherichia coli* MG1655 | V8I | Unknown | [3] |
| *Escherichia coli* MG1655 | Q38P | Reduced or diminished $IIA^{Glc}$-mediated allosteric inhibition | [4] |
| *Escherichia coli* MG1655 | V62L | Altered subunit interaction | [4] |
| *Escherichia coli* MG1655 | A68D | Altered subunit interaction | [3] |
| *Escherichia coli* MG1655 | D73A | Altered subunit interaction | [3] |
| *Escherichia coli* MG1655 | D73V | Altered subunit interaction | [3-4] |
| *Escherichia coli* MG1655 | G231D | Reduced or diminished sensitivity towards inhibitory effects of fructose 1,6-bisphosphate (FBP) | [3-4] |
| *Escherichia coli* MG1655 | G232S | Reduced or diminished sensitivity towards inhibitory effects of fructose 1,6-bisphosphate (FBP), altered interaction with phosphate ion | [3] |
| *Escherichia coli* MG1655 | I238T | Unknown | [3] |
| *Escherichia coli* MG1655 | M272I | Unknown | [4] |
| *Escherichia coli* MG1655 | M174I | Unknown | [3] |

* The start codon (encoding methionine) is numbered as 1st amino acid

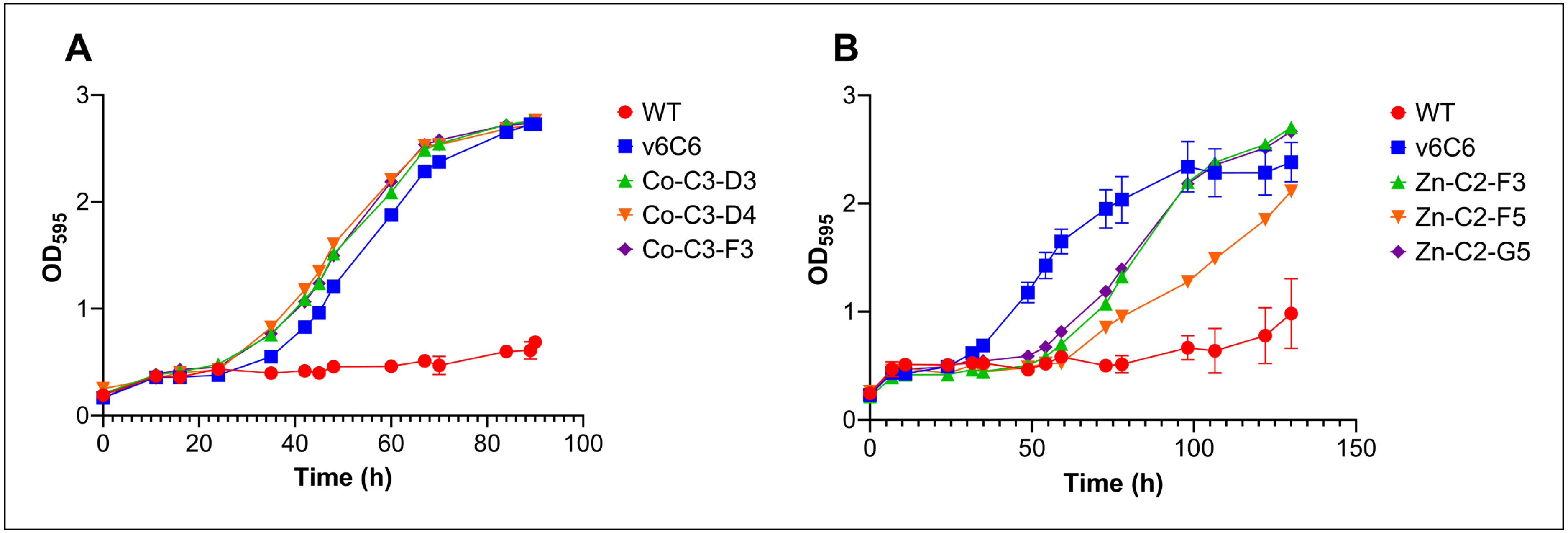


**Figure S1:** (**A**) The growth performance of variants Co-C3-D3 (green), Co-C3-D4 (orange), and Co-C3-F3 (purple) in 5% (v/v) glycerol was assessed in comparison to that of the WT (red) and v6C6 (blue) in a 96-well microplate. These variants were derived from a population that underwent 3 cycles of aALE. (**B**) The growth performance of variants Zn-C2-F3 (green), Zn-C2-F5 (orange), and Zn-C2-G5 (purple) in 5% (v/v) glycerol was assessed in comparison to that of the WT (red) and v6C6 (blue) in a 96-well microplate. These variants were derived from a population that underwent 2 cycles of aALE.

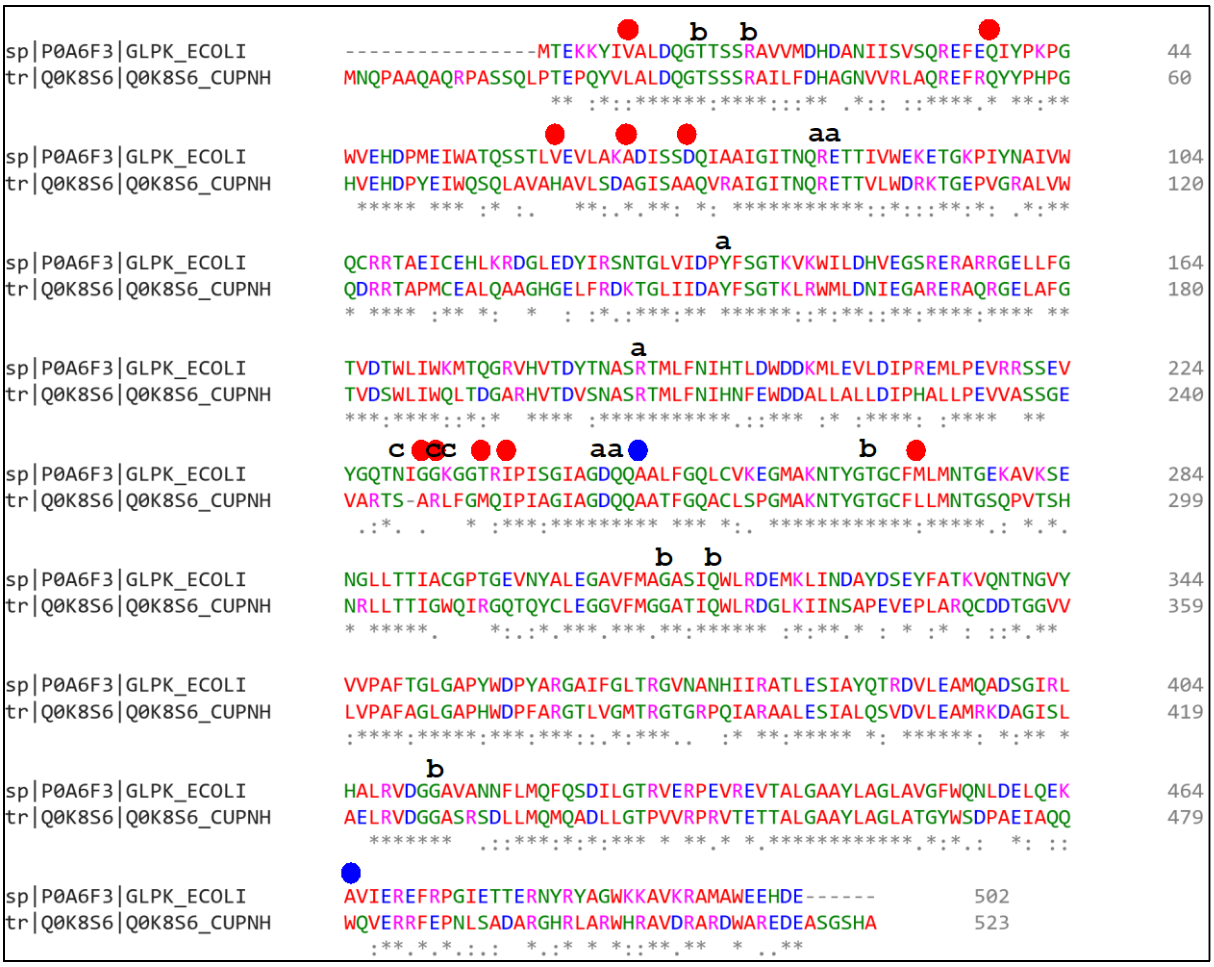


**Figure S2:** Protein sequence alignment between the glycerol kinase from *E. coli* (UniProt P0A6F3) and that from *C. necator* H16 (Q0K8S6) is illustrated. The alignment was performed using Clustal Omega [5]. Mutations discovered in glycerol-adapted *E. coli* and *C. necator* H16 through ALE are depicted with red dots and blue dots, respectively. Amino acids situated within the active site of the glycerol kinase are denoted using small letters, where 'a' indicates interaction with glycerol, 'b' denotes interaction with ADP, and 'c' represents interaction with a phosphate ion [6].

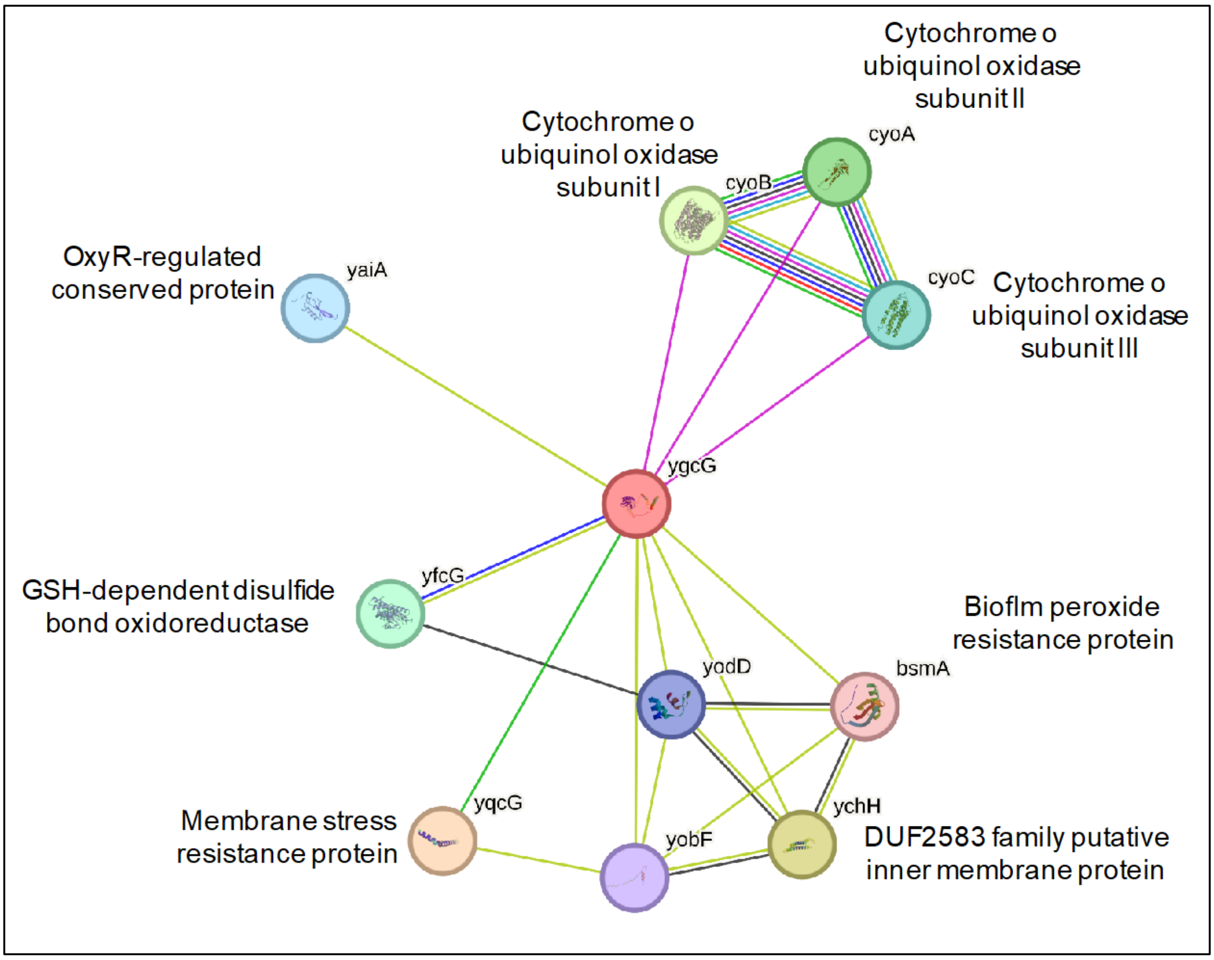


**Figure S3:** Functional protein association network of *E. coli* YgcG protein, created using STRING [7].
